# Gut microbiota modulation in response to combination of *Escherichia coli* Nissle 1917 and sugars: Lessons from comparative analysis of fecal microbiota of two healthy donors from 2019-2021

**DOI:** 10.1101/2022.06.10.495602

**Authors:** Debaleena Bhowmik, Kiran Heer, Manpreet Kaur, Saumya Raychaudhuri, Sandip Paul

## Abstract

The *Escherichia coli* Nissle 1917 strain (EcN) has shown its probiotic efficacy against many enteric pathogenic bacteria infecting human, including *Vibrio cholerae*, either alone or in combination with prebiotics. Understanding of these mechanisms of infection control requires the basic knowledge of probiotic mediated gut microbial community alterations especially in presence of different prebiotics. The present study has used the ex-vivo microbiota model and Next Generation Sequencing techniques to demonstrate the effect of EcN along with different sugars, namely glucose, galactose and starch, on the human gut microbiome community composition. The microbiome compositional changes have been observed at two different time-points, set one and a half years apart, in fecal slurries obtained from two donors. The study has indicated that the extent of microbiome alterations varies with different carbohydrate prebiotics and EcN probiotic and most of the alterations are broadly dependent upon the existing gut microbial community structure of the donors. The major distinct compositional changes have been found in the conditions where glucose and starch were administered, both with and without EcN, in spite of the inter-donor microbial community variation. Several of these microbiome component variations also remain consistent for both the time-points, including genus like *Bacteroides, Prevotella* and *Lactobacillus*. Altogether, the present study has shown the effectiveness of EcN along with glucose and starch towards specific changes of microbial community alterations independent of initial microbial composition. This type of model study can be implemented for hypothesis testing in case of therapeutic and prophylactic use of probiotic and prebiotic combinations.

## Introduction

*Escherichia coli* Nissle 1917 (abbreviated as EcN) was isolated by Prof. Alfred Nissle, German physician and microbiologist in 1917 during the World War I from a soldier who was resistant to a diarrheal outbreak (1, 2). Since its discovery more than 100 years ago, this particular strain has been studied intensely and also clinically employed in ameliorating ulcerative colitis (UC) symptoms (3). EcN is sold under the trade name mutaflor. Besides its therapeutic application in controlling burden of inflammatory bowel disease, EcN has also been exploited as a model organism to tackle other diseases as well. For example, genetically modified EcN producing cholera autoinducer 1 (CAI-1) is shown to restrict *Vibrio cholerae* virulence (4). EcN possess a robust iron uptake system as evidenced from genome analysis (5). Endowed with efficient iron uptake machinery, EcN has demonstrated its supremacy over *Salmonella typhimurium* and reduced the latter colonization by competing for iron in animal model (1). In another case, genetically engineered EcN can effectively sense and eliminate *Pseudomonas aeruginosa* in animal models (6). Not only tackling pathogens, EcN is also modified to convert ammonia to arginine as evidenced in mice model and human volunteers, thereby showing a great promise in clinical management of hyperammonemia (7).

Our engagement with EcN was started recently when we observed its efficacy in restricting the growth of *V. cholerae* in the presence of glucose in a co-culture growth condition (8). The *in vitro* observation was recapitulated in zebrafish infection model where *V. cholerae* colonization in the fish gut was drastically reduced by the combination of EcN and sugar (9). The cumulative outcome of these studies further raised the possibility of developing a combination therapy of glucose and EcN to treat cholera. Keeping view of the fact, we wanted to investigate the impact of the administration of glucose and EcN either alone or in combination on the community structure of gut microbiome or colonic microbiota. In addition to glucose, we also employed galactose and starch alone and also in combination with EcN to examine their impact on the gut microbiota. The fecal slurry ex-vivo microbiota model was adopted from fecal samples of two donors and the study was conducted in two phases where the changes in microbial composition were monitored with different sugar combinations along with EcN. The second set of replicates were setup after almost one and half years focusing mainly on the changes associated with the combined effect of glucose, starch and EcN addition on microbial community. The basis of this selection was our initial data that depicted most of the changes in the microbial community occurred in these particular sugars and EcN combination. With the help of metagenomics approach, we even could specifically show that changes in many of the microbial genus like *Bifidobacterium, Bacteroides* and *Lactobacillus* mostly occurred in this condition in a consistent way. Very few microorganisms showed changes with the galactose and starch sugar combinations, and also the amount of change was too small for a follow-up study with this combination. Besides, we also observed certain dramatic changes in the overall microbiome signature in almost one and half years at the donor level, especially reduction in the phylum *Actinobacteria* and an increase in *Proteobacteria*. Collectively, the present study highlighted the changes brought about by the exogenous administration of carbohydrates and probiotic on the structure of anaerobic microbial communities in fecal slurries obtained from two donors; of course, the changes on the community structure is also dependent on the existing diversity of the host gut microbiome.

## Results

### Inter-personal microbial community variation

We embarked on this study to evaluate the impact of the combination of probiotic *E. coli* Nissle 1917 (EcN) with various sugars on human gut microbiota. To achieve this feat, human fecal slurry based ex-vivo microbiota model was considered. We used similar sample characteristics, and gender but with different age of two donors to see the robustness of the probiotic, prebiotic effect. Also, different time point was chosen to prove the consistency of the changes. In the first phase, the inter-person variation in microbiome composition can be seen from the **Figure 1** irrespective of variation due to the addition of EcN and different sugars as prebiotics. We performed the statistical analysis including diversity indices and differential abundant genera for the samples from two individuals. The alpha diversity indices representing the within sample variation does not show a significant change (p>0.05) between two individuals **(Figure 2a)**. On the other hand, the beta diversity shows distinct clustering of samples between two donors asserting the fact that there is difference in microbiota composition, though statistically it wasn’t significant (p>0.05) **(Figure 2b)**. The difference in microbial composition between the donors can be seen from the Figure 1, the donor 1 had an overall higher abundance of *Bacteroides* as compared to donor 2; on the other hand, donor 2 had a dominant *Prevotella* population than donor 1. Other distinguishing genus between the two donors include *Megamonas, Collinsella, Catenibacterium* and *Faecalibacterium*.

**Figure 1.**
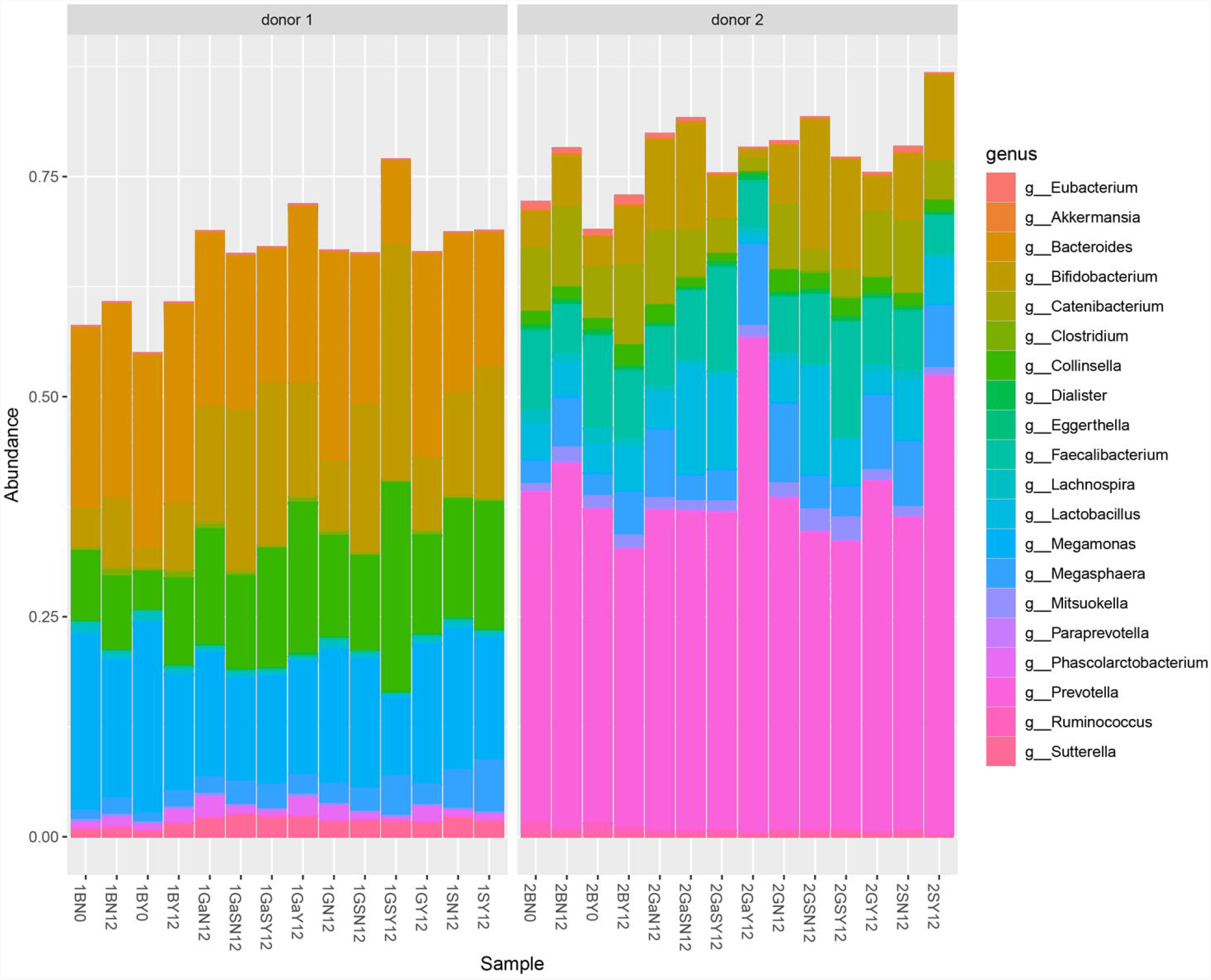
Taxonomic bar plot of the relative abundance of the top 20 genus in both donors accounting for more than 50% of the total microbial population, of phase 1 of the study.

**Figure 2.**
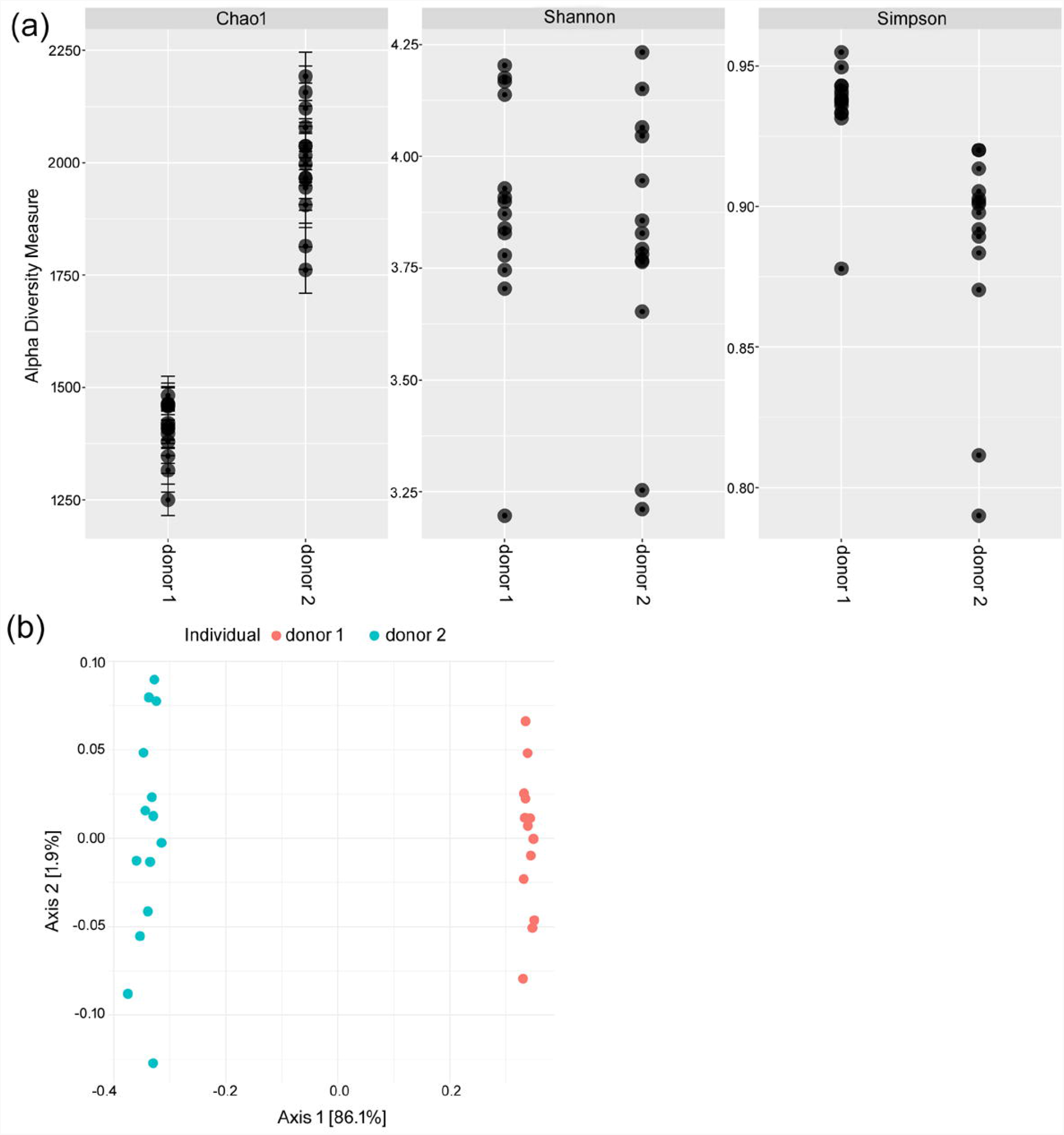
**(a)** Alpha diversity indices of the two donor samples. (**b)** Beta diversity plot of the two donor samples.

At the phylum level, **Figure 3**, the taxonomic distribution clearly depicts that donor 1 has a higher relative abundance of the phylum *Actinobacteria* as compared to donor 2, while phylum *Bacteroidetes*, is apparently more abundant in donor 2, and this difference between the donors, is highly significant, for both the phylum **(Supplementary Table 1)**. The *Firmicutes*:*Bacteroidetes* (F:B) ratio is crucial for a healthy gut microbiome and it is found that a higher F:B ratio is associated with obesity (11) and an increase in *Firmicutes* population also increases insulin resistance and other factors that may aid in T2DM development (12). As given in S**upplementary Table 2**, in all the blank condition samples from donor 2, the F:B ratio does not differ much, even in the presence of glucose+starch+EcN. In contrast to that, the F:B ratio of donor 1 was much higher compared to donor 2, and which increased further when glucose+starch+EcN was added. This outcome might be consistent with the fact that donor 1 was diabetic (**Table 1**). Along with that it was also observed that the difference in F:B ratio between the two donors were highly significant in phase 1 of the study (Paired T-test, p<0.001).

**Table 1.** Metadata containing all the details of both the donors.

| Criteria | Donor 1 | Donor 2 |
| --- | --- | --- |
| Gender | Male | Male |
| Age | 52 | 28 |
| Diet | Non vegetarian* | Mostly North Indian Vegetarian diet, Occasionally Non-veg (Once or twice a week). |
| Antibiotics usage 3 months prior to sample collection | No | No |
| Medicine usage | Metformin daily | No |
| Smoking | No | No |
| Alcohol usage | Last time December 2019 | Once/twice a month |
| Comorbidity | Diabetes, Asthma | No known comorbidity |
| Date of sample collection (1st time) | 23.07.19 | 11.09.19 |
| Date of sample collection (2nd time) | 01.03.21 | 01.03.21 |

\*a fiber-rich diet was adapted after the collection of first set of samples, with inclusion of plant-based fibers

**Figure 3.**
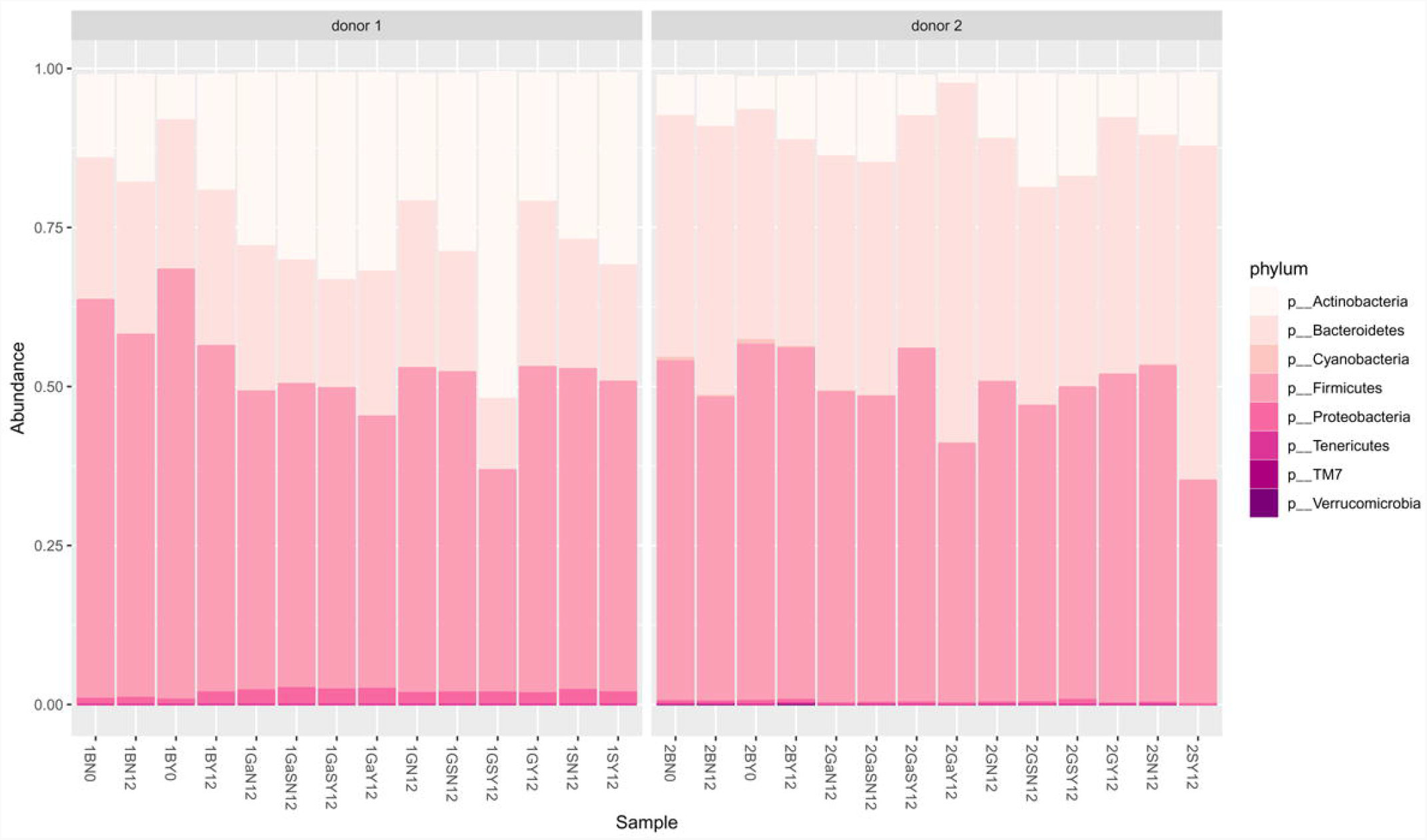
Taxonomic bar plot of the relative abundance of the different phylum in both donors from phase 1 of the study.

### Microbial community alterations due to factors like EcN and different sugars

We looked for the any changes associated with the addition of sugars (alone or in combination), EcN and sugar-EcN combinations in general. The alpha diversity analysis does not show much change in the microbial diversity for these three groups nor does the beta diversity analysis (data not shown). We found that the glucose+starch combination with EcN showing a consistent trend of microbial abundance change for several genera for both the individuals **(Supplementary Figure 1)**. In the first phase samples, for both the donors, *Bifidobacterium* showed more than 2-fold increase in the glucose+starch condition as compared to the blank and this increase was even more when incubated in the presence of EcN for donor 1 **(Supplementary Figure 1a)**. In case of *Megasphaera*, this pattern was observed only in donor 1, with almost 2.5 fold increase on addition of EcN **(Supplementary Figure 1d)**. Again, the reverse trend was observed in case of *Lachnospira*, almost 2 fold decrease on addition of EcN, for both donors **(Supplementary Figure 1e)**, while for *Blautia* this trend is only maintained in donor 1 **(Supplementary Figure 1b)**. The initial abundance of *Bifidobacterium* and further increase after treatment with sugar and EcN can be correlated with starch based food habits as starch and starch hydrolysates are preferred carbon sources for *Bifidobacteria* (13). A staple Indian food primarily constituted with rice and rice belongs to type 2 resistant starch (RS2). As evidenced, routine consumption of RS2 alters the abundance of some intestinal bacterial genera and species including *Bifidobacterium* species (14, 15). Interestingly, consumption of RS2 also decreases the proportion of *Lachnospiracea* and *Blautia* (15). This was also evidenced in our samples before and after treatment with sugar and EcN. Other than starch based diet, small molecules produced by *Bacillus* species also shown to promote growth of several strains of *Bifidobacterium* (16), further indicating cross feeding of metabolites modulate growth of gut anaerobes. It is therefore conceivable that exogenous administration of EcN may modulate metabolite production and thus control the growth of colonic anaerobic population in fecal slurry model system.

### Sustainable trends in microbial variation with effect from EcN and glucose, starch combination

Over the course of almost one and half years there was a remarkable decrease in the phylum *Actinobacteria*, while an increase in the phylum *Proteobacteria*, in both the donors, and which is quite evident from **Figure 4a**. As far as the F:B ratio was concerned, donor 2 did not show much of an alteration while in case of donor 1, there was a remarkable reduction in the F:B ratio from that of the earlier samples and which also remained stable even with the addition of glucose+starch+EcN **(Supplementary Table 2)**. This observation is very crucial based upon two major points – firstly, increased consumption of plant fiber and fiber-rich food contributes in stabilizing the F:B ratio in the gut and in this case donor 1 changed to a fiber-rich diet during the two years (**Table 1**); secondly, with a much stable F:B ratio, the gut is more resistant to changes.

**Figure 4.**
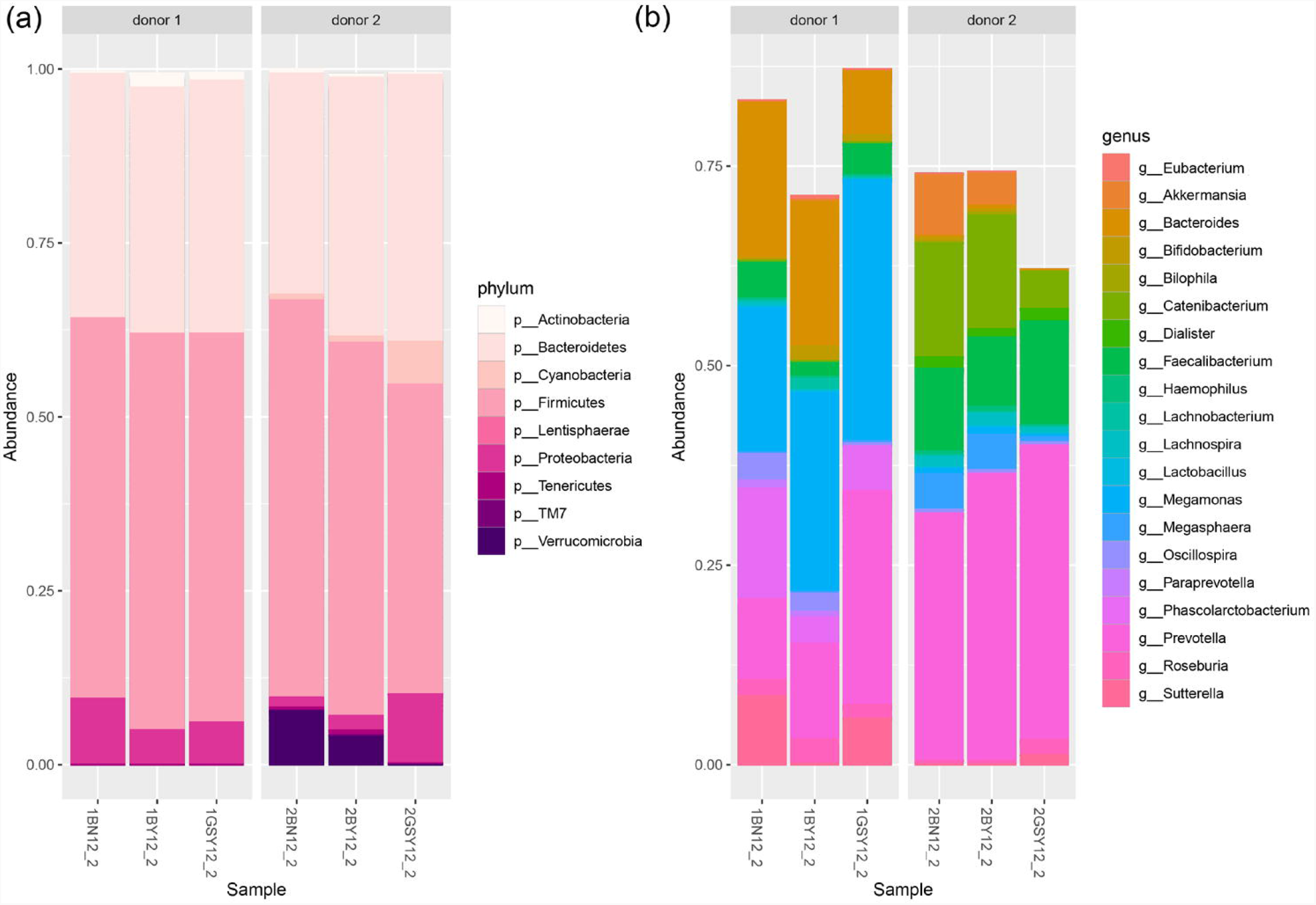
**(a)** Taxonomic bar plot of the relative abundance of the different phylum, for the phase 2 samples. **(b)** Taxonomic bar plot of the relative abundance of the different genus, for the phase 2 samples.

Based on the rarefied OTU table the Bray-Curtis distance for all the samples were calculated and hierarchically clustered, resulting in a dendogram as shown in **Figure 5**. The samples from the two different donors have appeared in separate clusters, signifying that there is quite a different microbial population between them.

**Figure 5.**
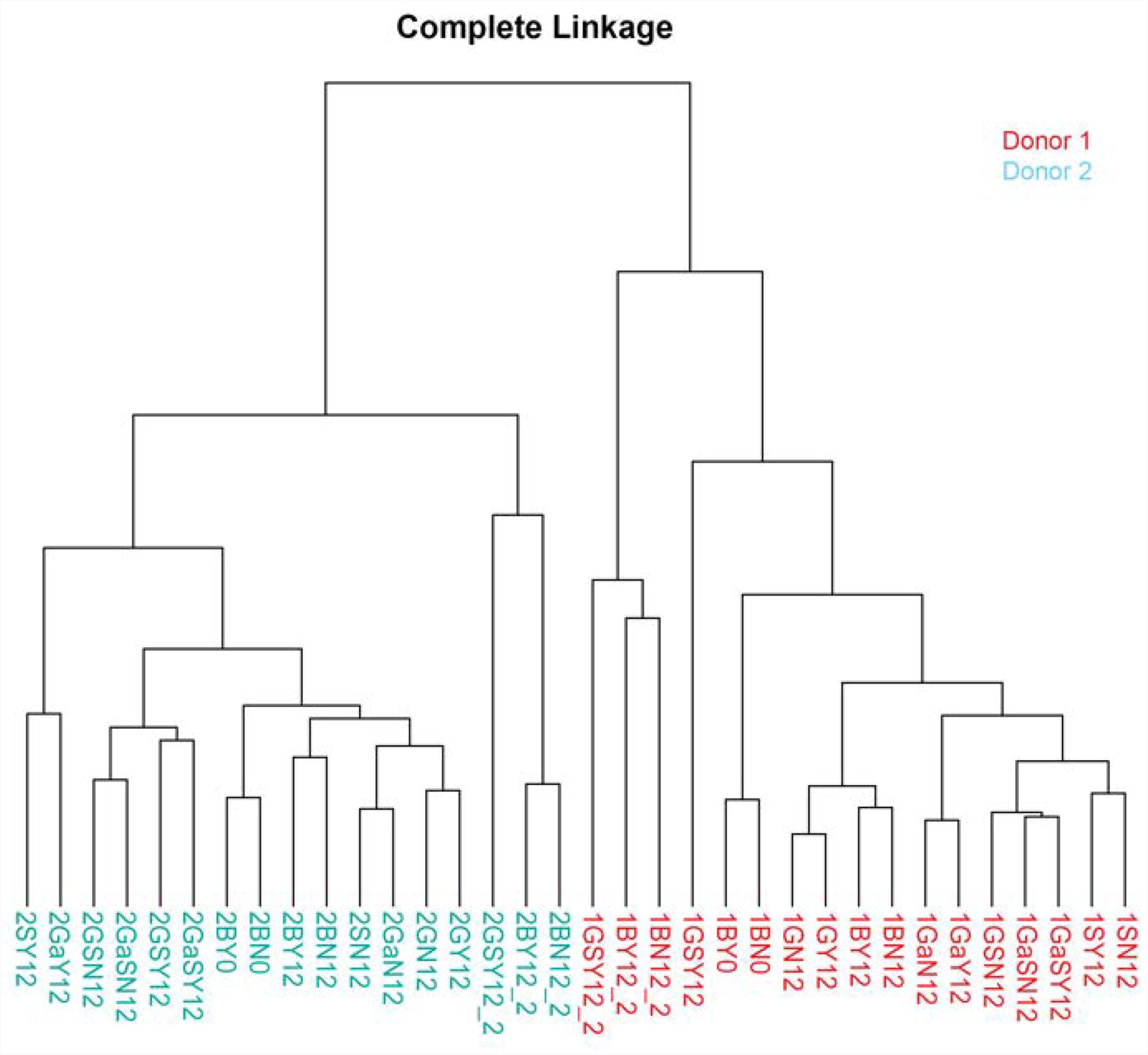
Dendogram formed by hierarchical clustering of the samples based on Bray-Curtis distance matrix, showing clear separation of samples from the two donors.

The second phase of collected samples showed a few changes in previous pattern at genus level in response to glucose+starch+EcN addition **(Figure 4b)**. The overall population of *Bifidobacterium* was reduced in both donors and also didn’t show any major change in the glucose+starch condition **(Supplementary Figure 1a)**. This change in *Bifidobacterium* population and trend may be attributed to the lifestyle or environment changes in both the donors, and since the initial population was far less as compared to the first set of collected samples, it probably didn’t show the expected changes with EcN addition when glucose+starch was added. In case of *Megasphaera*, donor 1 almost was diminished in its population in phase 2, while in case of donor 2, the trend with respect to phase 1 was reversed **(Supplementary Figure 1d)**; and for *Blautia* donor 1 showed a reversed trend while donor 2 remained the same **(Supplementary Figure 1b)**. Only for *Lachnospira* the trend in both the donors and for both the time of sample collection, was retained (**Supplementary Figure 1e)**.

On the other hand, the sustained trend can be seen for *Bacteroides*, which showed a decrease in abundance in the glucose+starch+EcN condition and also maintaining this trend at the end of 2 years while for the rest of the prominent taxa including *Megamonas, Sutterella* and *Streptococcus*, the initially observed trend changed with respect to the above condition (**Supplementary Figure 1d, 1g, 1h)**. Another important aspect for donor 1 as already highlighted was the spectacular increase in the *Prevotella* population, which was only present in donor 2 during the first time of sampling. Now coming to donor 2, the trend for almost all the major genus remained the same, *Prevotella, Lactobacillus* and *Faecalibacterium* showing an increase in glucose+starch+EcN condition while *Ruminococcus* and *Catenibacterium* reduced (**Supplementary Figure 1i, 1j, 1k, 1l)**. In addition to that, few more taxa of bacteria emerged as part of the dominant group like *Akkermensia* and *Mitsuokella* **(Figure 6)**.

**Figure 6.**
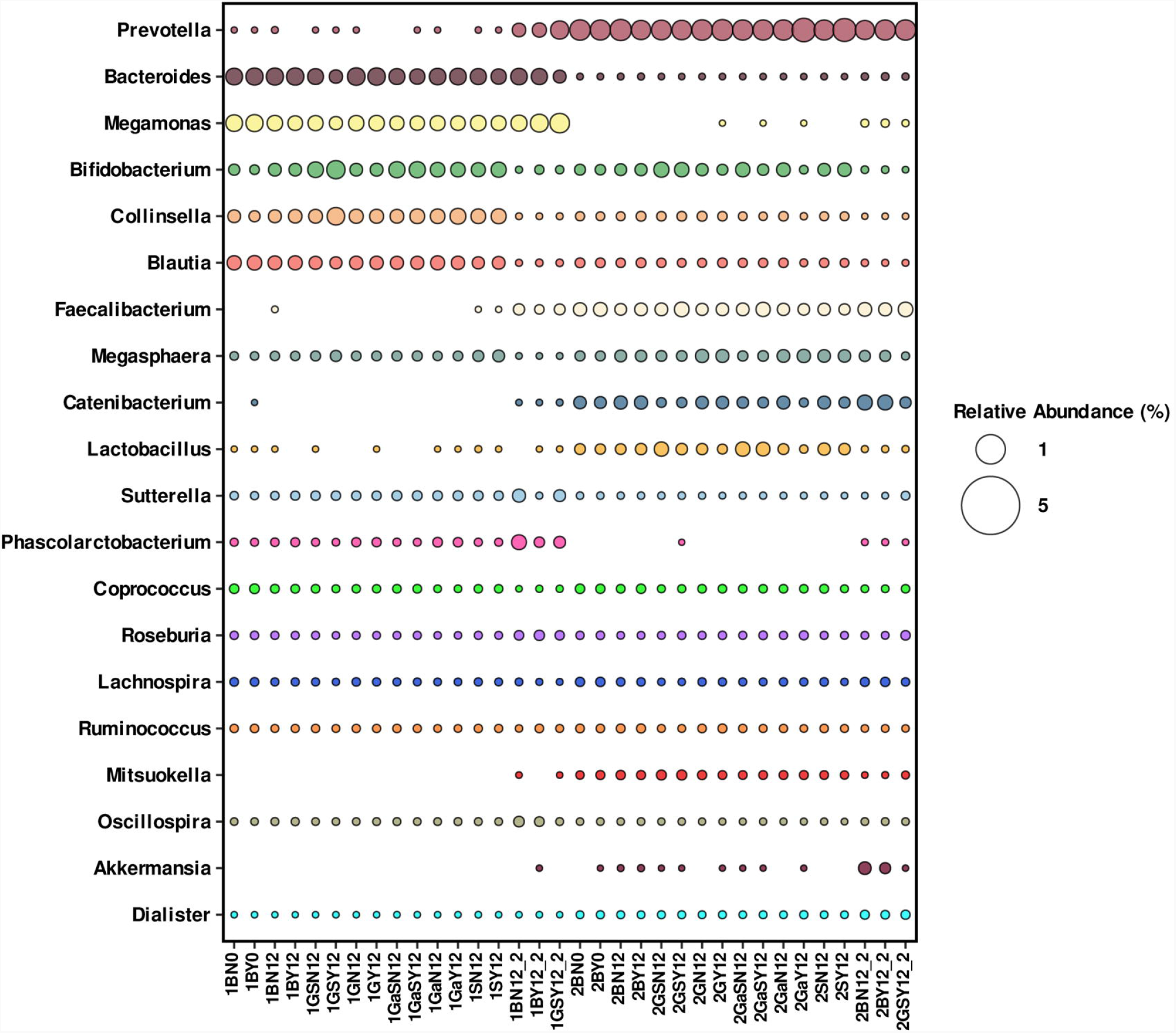
Bubble plot representation of genus-wise relative abundance of all the samples, highlighting those genus that showed most change between two phases and two donors.

Increase in *Akkermensia* is a known to be beneficial for the overall gut health (17) thus implying a beneficial transition for donor 2 during the second sample collection, but to check the actual effect of EcN on that would require further experimentation. It can also be concluded that the effectiveness of synbiotic (prebiotic + probiotic) depends largely on the initial microbiome composition, and the changes occurring might vary amongst individuals.

## Discussion

Previously, we demonstrated how diet combining glucose and commensal *E. coli* strains including probiotic *E. coli* Nissle 1917 (EcN) affect *V. cholerae* pathogenesis. Employing glucose transport mutant of *E. coli* that produces less acetate, it was concluded that acidic metabolites of glucose fermentation by *E. coli* strains reduce *V. cholerae* survival and colonization under in vitro co-culture growth and zebra fish infection model respectively (8, 9). Interestingly, accumulated evidences further suggested changes in gut microbial community by fermentable substrates and probiotic strains in various experimental models (18-20). It is therefore conceivable that *E. coli* and glucose combination which is shown to be effective in restricting *V. cholerae* pathogenesis may modulate the diversity of gut microbiota.

It is noteworthy to highlight that the glucose based oral rehydration solution recommended by WHO apparently increase fecal fluid loss (21, 22) and also enhances the production of virulence factors in *V. cholerae* (23). On the other hand, carbohydrates such as galactose and rice starch which are also known as prebiotics (24, 25) effectively reduced virulence expression in *V. cholerae* (23). Interestingly, *V. cholerae* is shown to adhere to starch granules (26), further bolstering the supremacy of starch-based oral rehydration therapy (ORT) over glucose based ORT. Moreover, starch is primarily fermented to produce beneficial short chain fatty acids (SCFA) in the colon, therefore, starch based ORT persists longer than glucose variety in the small intestine (27). Recently, the rice-based ORT was successfully field tested to treat cholera in Haiti (28).

Keeping this in mind, we wanted to investigate the effect of different sugars (prebiotics) alone and in combination with EcN (probiotic) on the structure of colonic microbiota in human fecal slurry ex-vivo microbiota model system. Though we demonstrated the efficacy of glucose and EcN combination in restricting *V. cholerae* growth and colonization in various experimental systems, we also kept starch in addition to glucose in our present study and our data clearly demonstrated glucose+starch+EcN combination worked better than glucose or starch or EcN alone in terms of promoting growth of certain beneficial anaerobic organisms and maintaining diversity of colonic microbiota in fecal slurry model. Some of the anaerobic organisms in the genera *Bacteroides, Prevotella, Faecalibacterium* etc. which are linked with the recovery phase of cholera infection (29) were also found to be maintained and promoted in the presence of glucose+starch+EcN combination.

In our study, we found that addition of sugars lowers the pH of the fecal slurry medium, especially in case of starch and also when combined with other sugars **(Supplementary Table 3)**. For the first set of samples from donor 1, a slight increase in *Bifidobacterium* was associated with the decrease in pH for both glucose+starch+EcN and starch+EcN conditions, while for the second set of samples, *Prevotella* increased in glucose+starch+EcN condition. For donor 2, *Prevotella* and *Faecalibacterium* increase was observed in glucose+starch+EcN condition alongside lowering of the pH, which remained consistent even after 2 years. *Prevotella* also retained a similar trend in the starch+EcN condition. It could be hypothesized that the decrease in pH due to addition of sugars and EcN facilitates the shift of microbiome to a healthier population but it would of course need a more data and also metabolite information.

Recently, Ansel Hsiao and colleagues have documented impact of interpersonal variation in microbiome on cholera pathogenesis (30). As evident, the microbiome with high taxonomic diversity having genera *Bacteroides, Clostridium, Blautia* etc. able to restrict colonization of *V. cholerae*. In contrast, the microbiome with low diversity and populated with *Streptococcus, Enterococcus faecalis* and *E. coli* referred as dysbiotic microbiome (such microbiome are common in cholera endemic areas) fail to restrict *V. cholerae* in infection in mice model (30). It would be interesting and challenging to examine the efficacy of glucose+starch+EcN combination in restricting *V. cholerae* infection in “dysbiotic microbiome” in animal and fecal slurry model. In the present study, we used equal amount of glucose and starch. As sugar has profound impact on gut microbiome, a range of varying concentrations of glucose and starch along with EcN should be evaluated. Additional studies are necessary to address these issues. As EcN is proven effective against other diarrheal pathogens, the combination of glucose+starch+EcN should be exploited against those pathogens as well in animal model to examine not only the restriction of pathogen but also restoration of gut microbiome. This warrants further investigation.

## Materials and Methods

### Ethical clearance, approval and metadata of the volunteers

Prior to the study, proper approval was obtained from the Institute’s Ethical Review Committee [IEC numbers are IEC (September2018) # 2 and IEC (January 2020) #1] and experiments were performed maintaining the established guidelines and regulations. Apart from that, written consents were obtained from the 2 donors of this study. A complete metadata was obtained from each volunteer in the form of a written questionnaire, recording their age, gender, food habit, antibiotic usage and or other medication, and health status, at the time of sample collection **(Table 1)**. The study was conducted in two phases. In the first phase, the effect of different sugars, alone and in combination, along with EcN on donors’ fecal microbial composition were monitored. In case of the second phase, almost after one and half years of first phase, microbial community alterations were studied with the combined effect of glucose, starch and EcN. A detail depiction of the study design with different probiotic and prebiotic combinations is given in **Supplementary Figure 2**. It should be noted that one of the donors took antibiotics and anti-protozoan drugs during the intermediate span of one and half years but not within 3 months before giving fecal samples. None of the volunteers suffered any form of diarrhea at the point of donating samples.

### Sample collection, DNA extraction and sequencing

Fecal samples from two donors were freshly collected in a sterile container. Samples were processed immediately by preparing stock of 20% slurry (w/v) in 50mM PBS buffer following published protocol (31, 32). Slurries were diluted to 10% in 50mM PBS. 50ml of 10% slurries were distributed in serum vial and mixed with exponentially grown 5X 10e6 EcN and 1% sugars (glucose/galactose/rice-starch; 1% each) alone and in combination of EcN and each sugar. Serum vials were purged with anaerobic gas mixtures, followed by sealing and incubated for 12 hr at 37C with agitation at 150 rpm. After stipulated period of incubation, slurries were collected and shipped to Medgenome Labs Ltd under cold condition for further processing. Total fecal DNA isolation, library preparation and next generation sequencing were performed at Medgenome. Bioinformatics data analysis was done at our end. The details of the donors are given in Table 1, while that of the different conditions and treatment of the samples are given in **Supplementary Table 3**.

### Construction of *Escherichia coli* Nissle 1917 (EcN) rifampicin variant

Sub-culturing of EcN was done in increasing concentrations of rifampicin to create a rifampicin-resistant mutant. Passaging of bacterial culture was done starting from 5µg/ml upto 75µg/ml of rifampicin (31). EcN/Rif mutant was then maintained on LB/Rif (75µg/ml) plates and further confirmed by sequencing. In order to check the survival of EcN/Rif, after 12 hrs. of incubation of fecal slurry with EcN/Rif, 1ml of fecal slurry was collected from serum bottle and dilutions were made upto 10^-6 and then spotting was done on LB/Rif (75ug/ml) plates.

### Analysis of 16S rDNA sequencing data

The V3-V4 region of the 16S rDNA was sequenced and the quality of the raw reads were checked using FastQC (33). Further processing was performed with standard QIIME1 pipeline (34) which includes, primer trimming, filtering and OTU (operational taxonomic unit) picking. Closed reference OTU picking with 97% sequence similarity against Greengenes database (version 13.8, 99% OTU cluster) (10) was used, assigning the taxonomy at confidence 0.8. The assigned taxonomies were summarized into relative abundance tables at both phylum and genus level for further taxonomic analysis.

### Sample characteristics and sequence statistics

The paired-end amplicon sequencing of the V3-V4 region of 16S rRNAyielded a total of 30,914,413 reads for all the 34 samples and after merging the reads, primer trimming, and filtering based on overall quality (read length and error rate), a total of 7,104,596 filtered reads was obtained **(Supplementary Table 4)**. These reads were then used for closed reference OTU picking based on Greengenes database (version 13.8) (10) and it resulted in 6,514 OTUs for all the samples. Relative abundance table at genus level classification yielded a total of 179 taxa, though not all were classified till the genus level. We checked the top 20 taxa based on their mean relative abundance genus level classification and found that they accounted for >50% of the entire population for each sample.

### Representation of the taxonomic data and clustering

For the differential abundance, alpha diversity and beta diversity analysis and significance testing, phyloseq R package (35) was used. For beta diversity, the Bray-Curtis distance matrix was used with MDS method. The bubble plot for genus level taxa representation was constructed using R packages ggplot2 (36) and reshape2 (37). For the dendogram, the Bray-Curtis distance was used and the samples were clustered using the “complete” method of hierarchical clustering; for this the R packages bcdist and hclust were used.

## Author contributions

SR conceived the idea and designed experiments with MK, DB, KH and SP; KH and MK collected fecal samples from donors, processed and performed fecal slurry experiments with EcN and sugars; DB and SP processed and analyzed microbiome data.SR, SP and DB wrote the manuscript; all authors gave insightful comments and approved the manuscript.

## Data availability

The raw read sequences are available at NCBI SRA database under the BioProject Accession ID PRJNA816292.

## Acknowledgement

We gratefully acknowledge all donors for fecal samples. We extend our appreciation to all donors for their support. SR extendsgratitude to lab membersof SR, for critical evaluation of the entire manuscript. DB thanks Abhishake Lahiri of her lab for his technical help and manuscript evaluation.

## Supplementary Material

**Supplementary Table 1** – Statistical significance of all phylum between the two donors.

**Supplementary Table 2** – F:B ratios of the samples from both the donors during both the phases, and their statistical significance testing. The highlighted samples are the ones whose conditions were replicated in phase 2 of the study.

**Supplementary Table 3** – Details of all the different conditions and treatments of the samples.

**Supplementary Table 4** – Sample-wise counts of the raw reads and filtered reads (Filter method is described in the method section)

**Supplementary Figure 1** – Graphical representation of the changes in genus through different treatments (sugar and EcN addition) and in different donors in both the phases of the study. The genus which either retained their trend or altered it remarkably, were selected

**Supplementary Figure 2** – Diagrammatic representation of all the combinations of sugars used for the samples from two donors at two-time intervals.

