## Supplementary figures and images for "Gut microbiota modulation in response to combination of *Escherichia coli* Nissle 1917 and sugars: Lessons from comparative analysis of fecal microbiota of two healthy donors from 2019-2021"

### Supplementary Figure 1

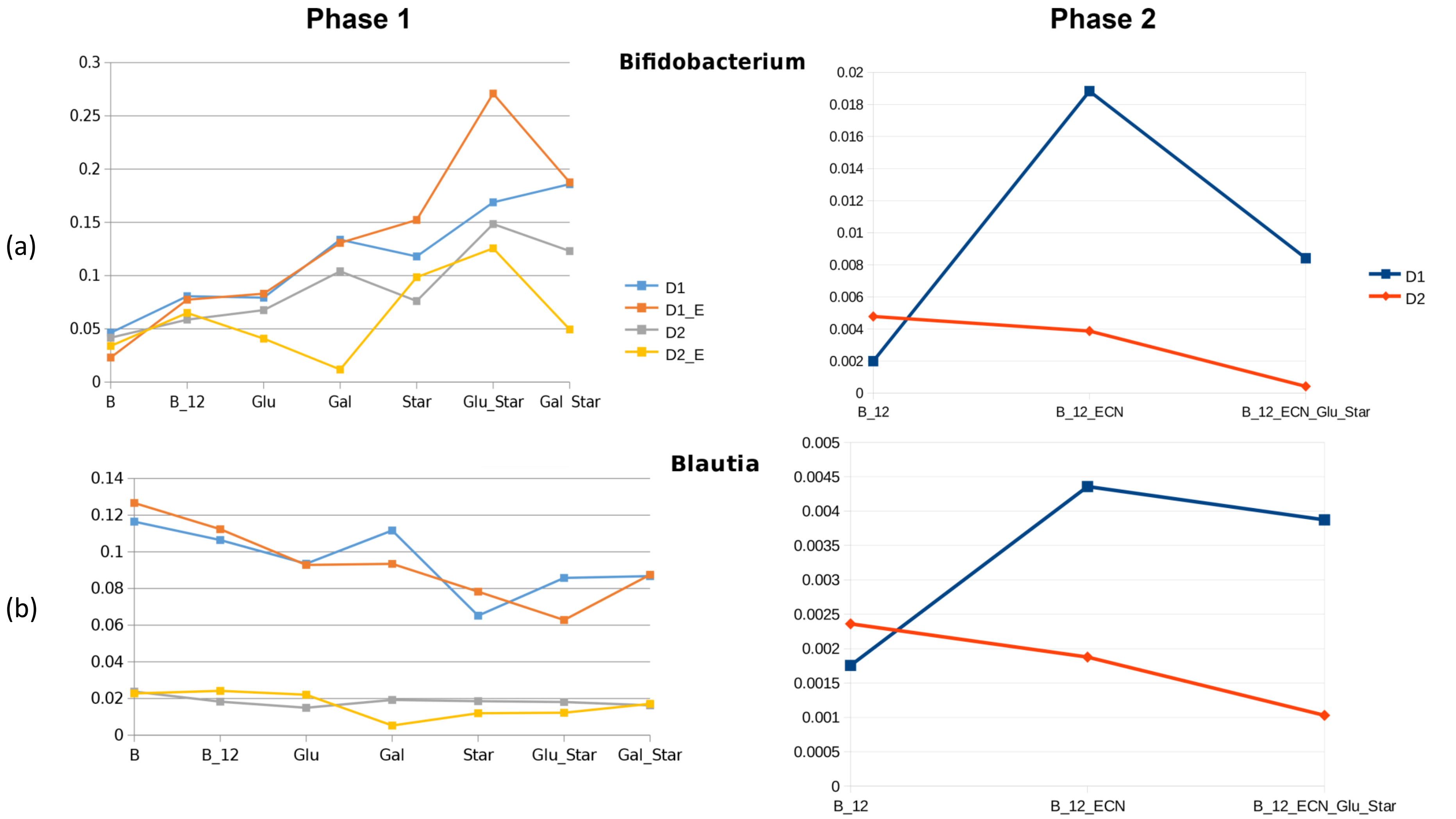

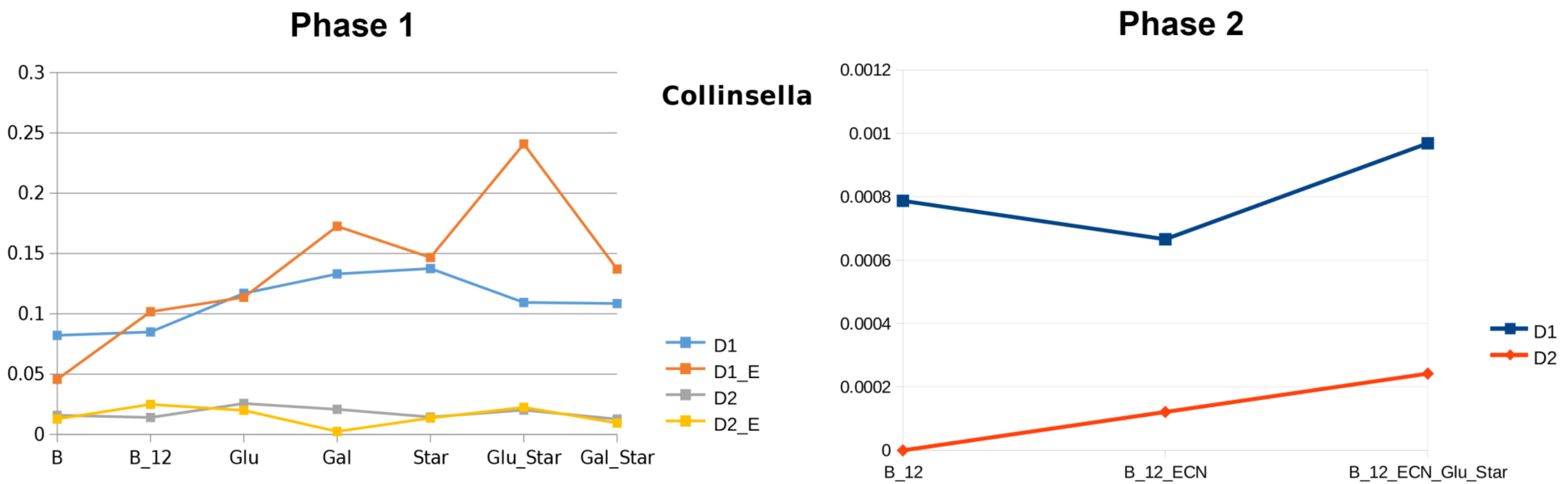

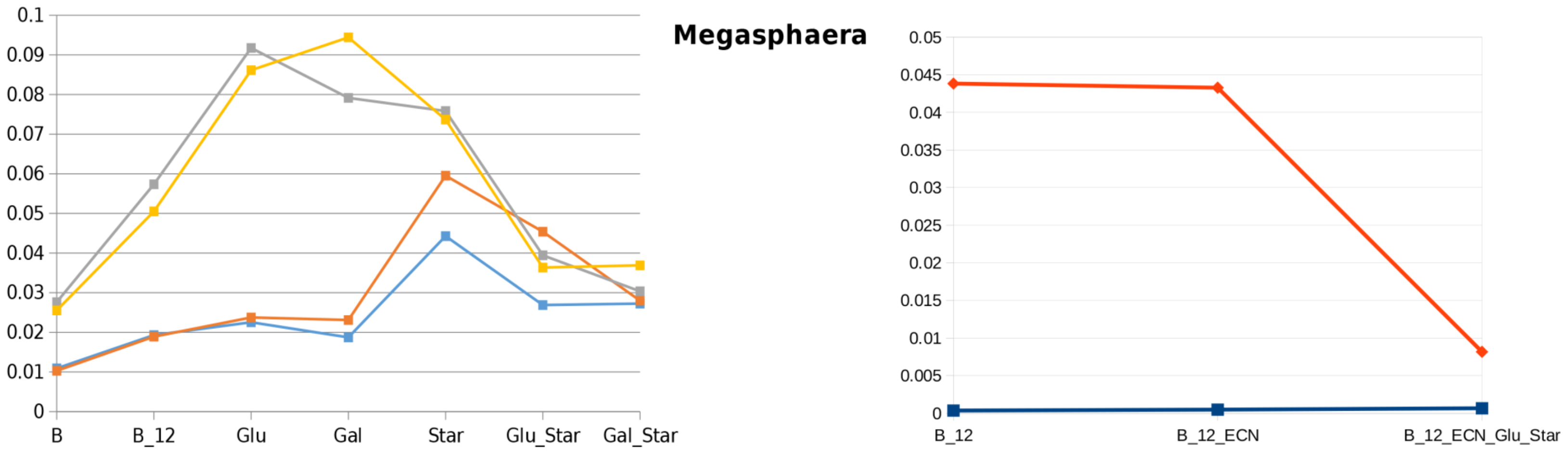

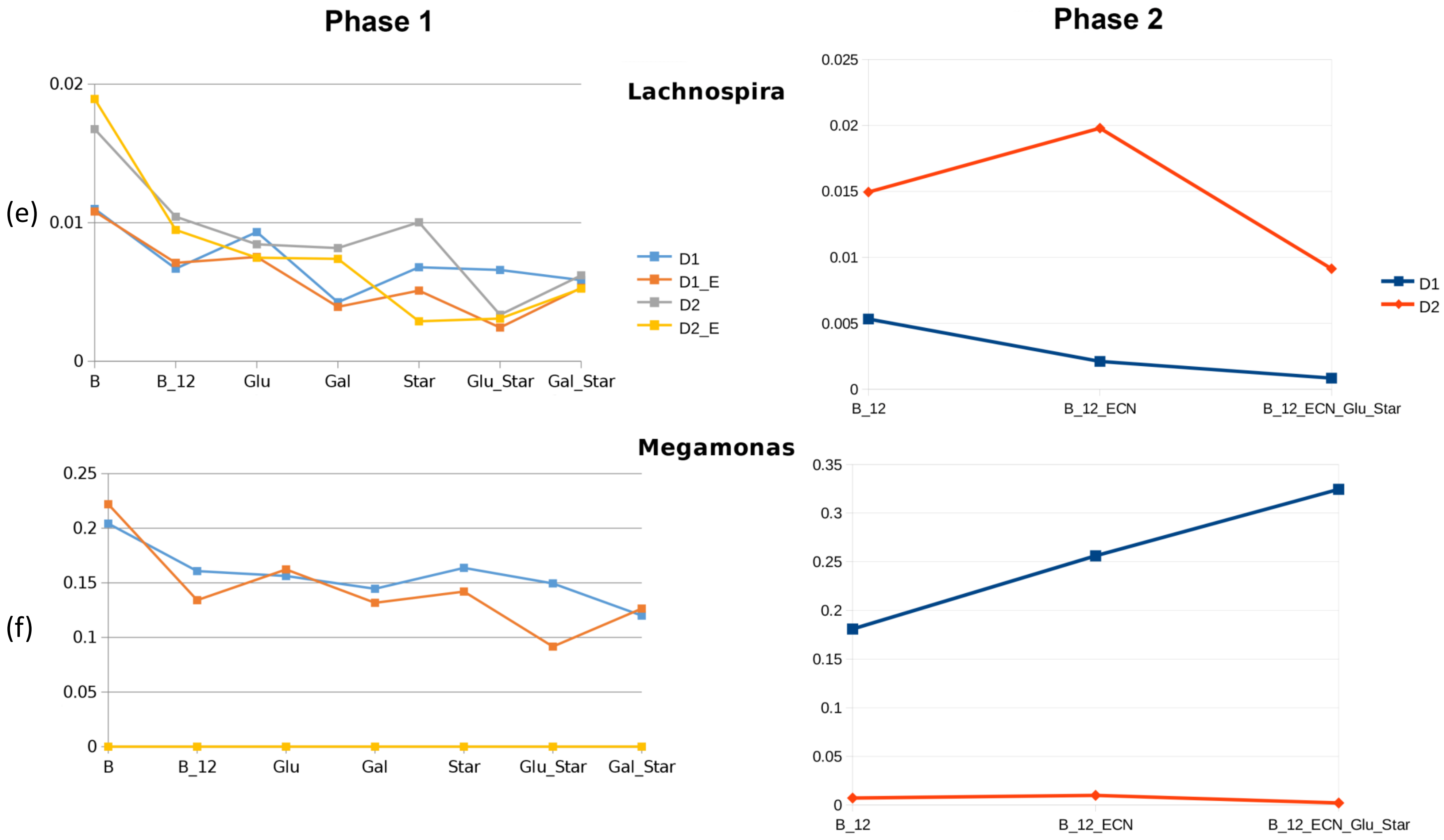

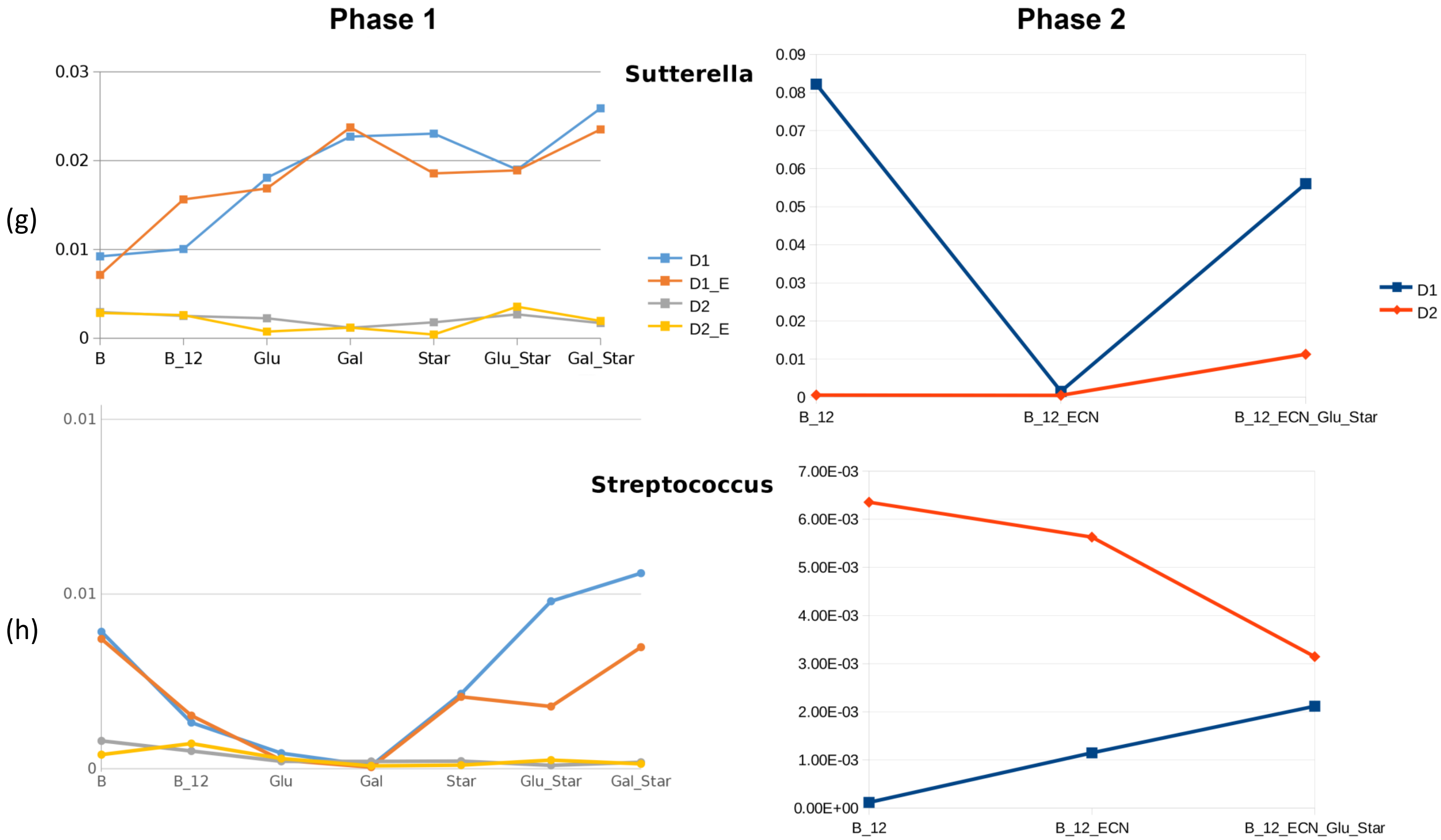

**Phase 1** **Phase 2**

**Faecalibacterium**

(i)

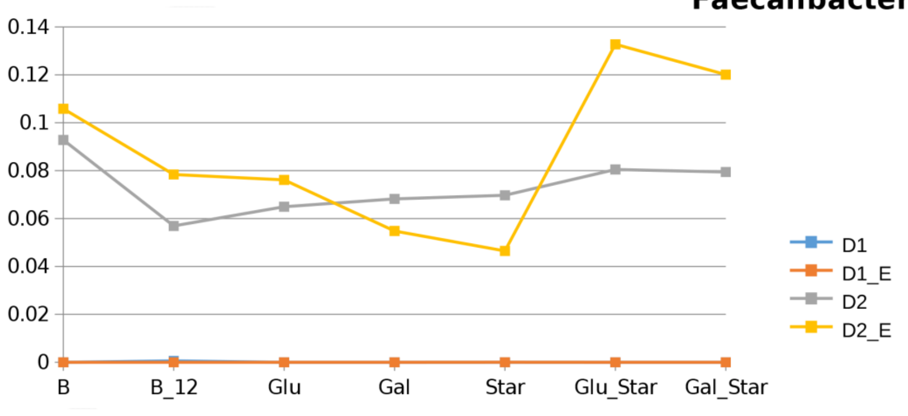


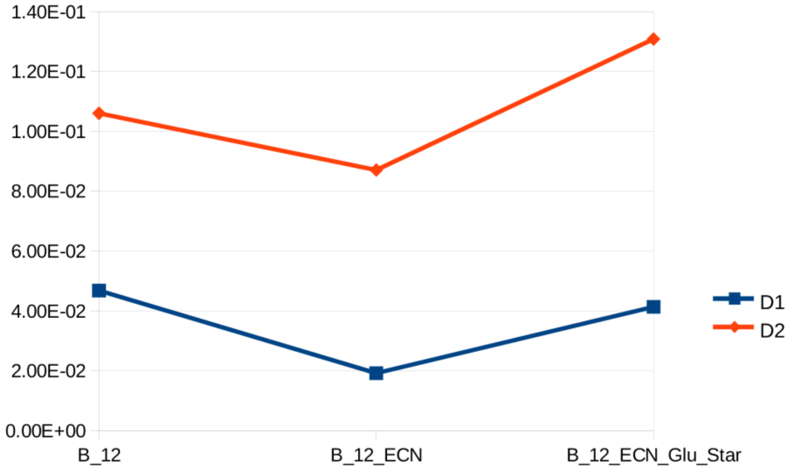


**Ruminococcus**

(j)

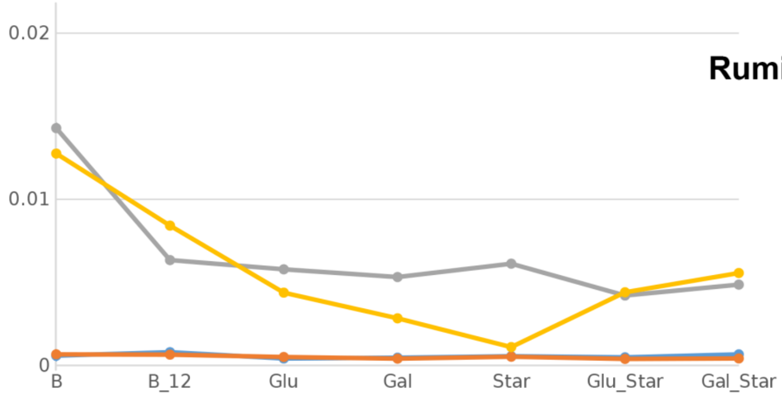


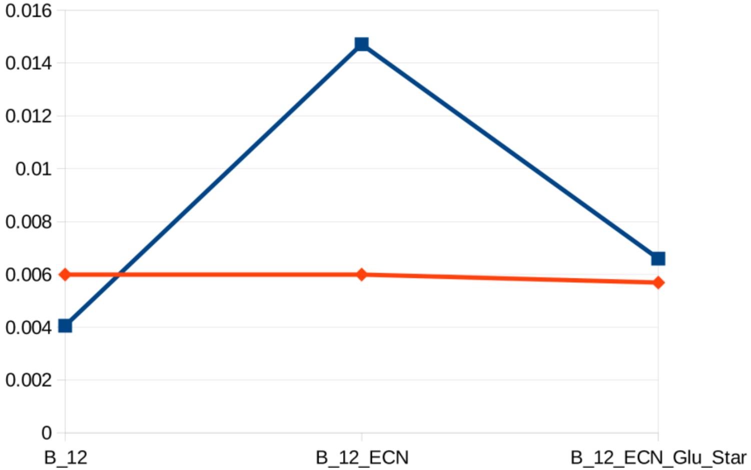

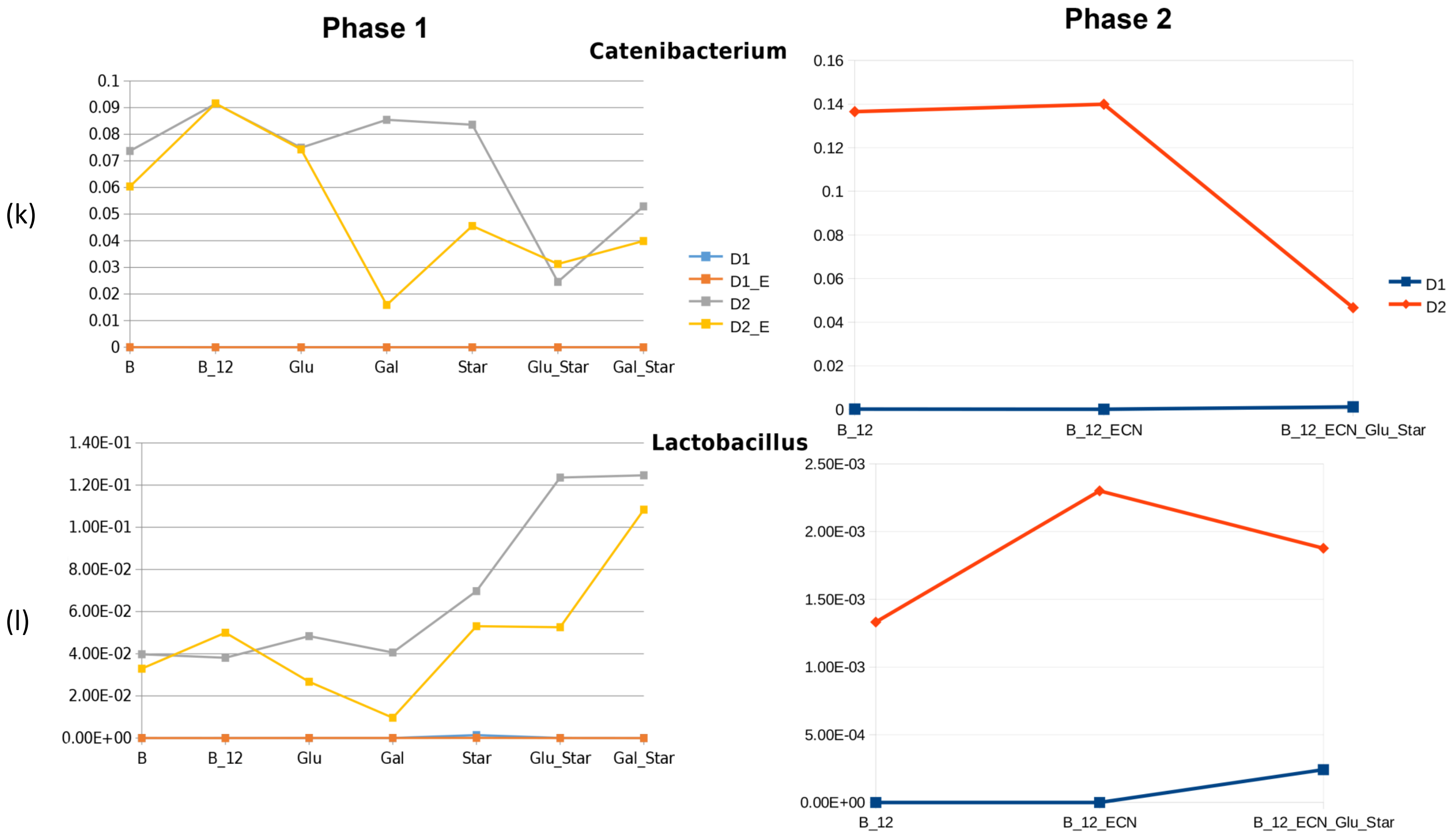
