## Supplementary Figure 2 for "Gut microbiota modulation in response to combination of *Escherichia coli* Nissle 1917 and sugars: Lessons from comparative analysis of fecal microbiota of two healthy donors from 2019-2021"

Phase one sample collection

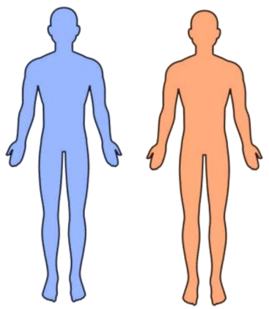

Phase two sample collection

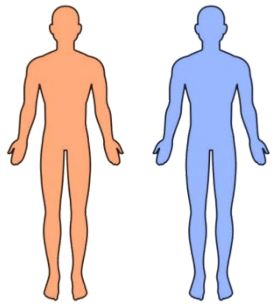

537 days

587 days

Fecal samples

Fecal slurry

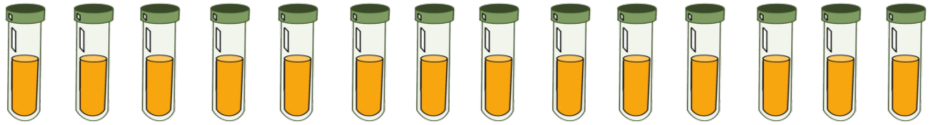

| Time (hrs) | 0 | 0 | 12 | 12 | 12 | 12 | 12 | 12 | 12 | 12 | 12 | 12 | 12 | 12 |
|---|---|---|---|---|---|---|---|---|---|---|---|---|---|---|
| **EcN** | X | ✓ | X | ✓ | X | ✓ | X | ✓ | X | ✓ | X | ✓ | X | ✓ |
| **Glucose** | X | X | X | X | ✓ | ✓ | X | X | X | X | ✓ | ✓ | X | X |
| **Galactose** | X | X | X | X | X | X | ✓ | ✓ | X | X | X | X | ✓ | ✓ |
| **Starch** | X | X | X | X | X | X | X | X | ✓ | ✓ | ✓ | ✓ | ✓ | ✓ |
| Sample Names | 1BN0/2BN0 | 1BY0/2BY0 | 1BN12/2BN12 | 1BY12/2BY12 | 1GN12/2GN12 | 1GY12/2GY12 | 1GaN12/2GaN12 | 1GaY12/2GaY12 | 1SN12/2SN12 | 1SY12/2SY12 | 1GSN12/2GSN12 | 1GSY12/2GSY12 | 1GaSN12/2GaSN12 | 1GaSY12/2GaSY12 |

V3-V4 amplicon sequencing and data analysis

Fecal samples

Fecal slurry

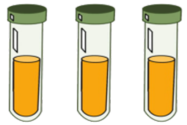

| Time (hrs) | 12 | 12 | 12 |
|---|---|---|---|
| **EcN** | X | ✓ | ✓ |
| **Glucose** | X | X | ✓ |
| **Starch** | X | X | ✓ |
| Sample Names | 1BN12_2/2BN12_2 | 1BY12_2/2BY12_2 | 1GSY12_2/2GSY12_2 |
