## Supplementary Table 1 for "Gut microbiota modulation in response to combination of *Escherichia coli* Nissle 1917 and sugars: Lessons from comparative analysis of fecal microbiota of two healthy donors from 2019-2021"

| **phylum** | **raw p** | **adjp** | **fdr** |
|---|---|---|---|
| p__Proteobacteria | 0 | 0 | 0.0012857 |
| p__Bacteroidetes | 0 | 0 | 0.0012857 |
| p__TM7 | 0 | 0 | 0.0012857 |
| p__Actinobacteria | 0 | 0 | 0.0012857 |
| p__Cyanobacteria | 0 | 0 | 0.0012857 |
| p__Tenericutes | 0 | 0 | 0.0012857 |
| p__Verrucomicrobia | 0 | 0 | 0.0012857 |
| p__Firmicutes | 0.79 | 0.89 | 0.817 |
| p__Fusobacteria | 0.82 | 0.89 | 0.817 |

**Supplementary Table 1** – Statistical significance of all phylum between the two donors.
