## Supplementary Table 2 for "Gut microbiota modulation in response to combination of *Escherichia coli* Nissle 1917 and sugars: Lessons from comparative analysis of fecal microbiota of two healthy donors from 2019-2021"

| Sample_ID | Phase | Donor | Firmicutes:Bacteroidetes ratio |
|---|---|---|---|
| 1BN0 | 1 | 1 | 2.8429 |
| 1BN12 | 1 | 1 | 2.4064 |
| 1BY0 | 1 | 1 | 2.9001 |
| 1BY12 | 1 | 1 | 2.2454 |
| 1GSY12 | 1 | 1 | 3.1399 |
| 1GaN12 | 1 | 1 | 2.0681 |
| 1GaSN12 | 1 | 1 | 2.4785 |
| 1GaSY12 | 1 | 1 | 2.8079 |
| 1GaY12 | 1 | 1 | 1.8921 |
| 1GN12 | 1 | 1 | 1.9617 |
| 1GSN12 | 1 | 1 | 2.6837 |
| 1GY12 | 1 | 1 | 1.9848 |
| 1SN12 | 1 | 1 | 2.5018 |
| 1SY12 | 1 | 1 | 2.6818 |
| 2BN0 | 1 | 2 | 1.4279 |
| 2BN12 | 1 | 2 | 1.1502 |
| 2BY0 | 1 | 2 | 1.5777 |
| 2BY12 | 1 | 2 | 1.7287 |
| 2GSY12 | 1 | 2 | 1.5083 |
| 2GaN12 | 1 | 2 | 1.3414 |
| 2GaSN12 | 1 | 2 | 1.3296 |
| 2GaSY12 | 1 | 2 | 1.5470 |
| 2GaY12 | 1 | 2 | 0.7333 |
| 2GN12 | 1 | 2 | 1.3387 |
| 2GSN12 | 1 | 2 | 1.3766 |
| 2GY12 | 1 | 2 | 1.3053 |
| 2SN12 | 1 | 2 | 1.4856 |
| 2SY12 | 1 | 2 | 0.6783 |

**ANOVA**

| P-value | F critical |
|---|---|
| **3.58523E-09** | **4.225201273** |

**Paired t-test**

| | |
|---|---|
| P (T<=t) one-tail | **5.54342E-08** |
| t Critical one-tail | 1.770933396 |
| P (T<=t) two-tail | **1.10868E-07** |
| t Critical two-tail | 2.160368656 |

| Sample_ID | Phase | Donor | Firmicutes:Bacteroidetes ratio |
|---|---|---|---|
| 1BN12_2 | 2 | 1 | 1.5570 |
| 1BY12_2 | 2 | 1 | 1.6209 |
| 1GSY12_2 | 2 | 1 | 1.5449 |
| 2BN12_2 | 2 | 2 | 1.7969 |
| 2BY12_2 | 2 | 2 | 1.4552 |
| 2GSY12_2 | 2 | 2 | 1.1668 |

**Supplementary Table 2** – F:B ratios of the samples from both the donors during both the phases, and their statistical significance testing. The yellow highlighted samples are the ones whose conditions were replicated in phase 2 of the study. The top table represents data from phase 1 while that of phase 2 is represented in the bottom table.
