## Supplementary Table 3 for "Gut microbiota modulation in response to combination of *Escherichia coli* Nissle 1917 and sugars: Lessons from comparative analysis of fecal microbiota of two healthy donors from 2019-2021"

| Sample | Sample ID | Sugar | EcN | Incubation time (hrs) | pH | Incubation temp | Incubation condition | rif spotting test | Sugar amount | ECN amount | Sample collection set | Other conditions (if any) |
|---|---|---|---|---|---|---|---|---|---|---|---|---|
| FS001 | 1BN0 | NA | No | 0 | NA | NA | Anaerobic conditions | NA | NA | NA | 1 | 1.5mM cysteine |
| FS002 | 1BY0 | NA | Yes | 0 | NA | NA | Anaerobic conditions | Positive | NA | 5 X 10^6 cells/ml | 1 | 1.5mM cysteine |
| FS003 | 2BN0 | NA | No | 0 | NA | NA | Anaerobic conditions | NA | NA | NA | 1 | 1.5mM cysteine |
| FS004 | 2BY0 | NA | Yes | 0 | NA | NA | Anaerobic conditions | Positive | NA | 5 X 10^6 cells/ml | 1 | 1.5mM cysteine |
| FS005 | 1BN12 | NA | No | 12 | 5 to 6 | 37C | Anaerobic conditions | NA | NA | NA | 1 | 1.5mM cysteine |
| FS006 | 1BY12 | NA | Yes | 12 | 4 to 5 | 37C | Anaerobic conditions | Positive | NA | 5 X 10^6 cells/ml | 1 | 1.5mM cysteine |
| FS007 | 2BN12 | NA | No | 12 | 6 to 7 | 37C | Anaerobic conditions | NA | NA | NA | 1 | 1.5mM cysteine |
| FS008 | 2BY12 | NA | Yes | 12 | 7 to 8 | 37C | Anaerobic conditions | Positive | NA | 5 X 10^6 cells/ml | 1 | 1.5mM cysteine |
| FS009 | 1GSN12 | Glucose + Starch | No | 12 | 5 to 6 | 37C | Anaerobic conditions | NA | 1.0% | NA | 1 | 1.5mM cysteine |
| FS010 | 1GSY12 | Glucose + Starch | Yes | 12 | 4 to 5 | 37C | Anaerobic conditions | Positive | 1.0% | 5 X 10^6 cells/ml | 1 | 1.5mM cysteine |
| FS011 | 2GSN12 | Glucose + Starch | No | 12 | 4 to 5 | 37C | Anaerobic conditions | NA | 1.0% | NA | 1 | 1.5mM cysteine |
| FS012 | 2GSY12 | Glucose + Starch | Yes | 12 | 4 to 5 | 37C | Anaerobic conditions | Positive | 1.0% | 5 X 10^6 cells/ml | 1 | 1.5mM cysteine |
| FS013 | 1GN12 | Glucose | No | 12 | 5 to 6 | 37C | Anaerobic conditions | NA | 1.0% | NA | 1 | 1.5mM cysteine |
| FS014 | 1GY12 | Glucose | Yes | 12 | 4 to 5 | 37C | Anaerobic conditions | Positive | 1.0% | 5 X 10^6 cells/ml | 1 | 1.5mM cysteine |
| FS015 | 2GN12 | Glucose | No | 12 | 5 to 6 | 37C | Anaerobic conditions | NA | 1.0% | NA | 1 | 1.5mM cysteine |
| FS016 | 2GY12 | Glucose | Yes | 12 | 4 to 5 | 37C | Anaerobic conditions | Positive | 1.0% | 5 X 10^6 cells/ml | 1 | 1.5mM cysteine |
| FS017 | 1GaSN12 | Galactose + Starch | No | 12 | 5 to 6 | 37C | Anaerobic conditions | NA | 1.0% | NA | 1 | 1.5mM cysteine |
| FS018 | 1GaSY12 | Galactose + Starch | Yes | 12 | 4 to 5 | 37C | Anaerobic conditions | Positive | 1.0% | 5 X 10^6 cells/ml | 1 | 1.5mM cysteine |
| FS019 | 2GaSN12 | Galactose + Starch | No | 12 | 4 to 5 | 37C | Anaerobic conditions | NA | 1.0% | NA | 1 | 1.5mM cysteine |
| FS020 | 2GaSY12 | Galactose + Starch | Yes | 12 | 4 to 5 | 37C | Anaerobic conditions | Positive | 1.0% | 5 X 10^6 cells/ml | 1 | 1.5mM cysteine |
| FS021 | 1GaN12 | Galactose | No | 12 | 5 to 6 | 37C | Anaerobic conditions | NA | 1.0% | NA | 1 | 1.5mM cysteine |
| FS022 | 1GaY12 | Galactose | Yes | 12 | 4 to 5 | 37C | Anaerobic conditions | Positive | 1.0% | 5 X 10^6 cells/ml | 1 | 1.5mM cysteine |
| FS023 | 2GaN12 | Galactose | No | 12 | 5 to 6 | 37C | Anaerobic conditions | NA | 1.0% | NA | 1 | 1.5mM cysteine |
| FS024 | 2GaY12 | Galactose | Yes | 12 | 4 to 5 | 37C | Anaerobic conditions | Positive | 1.0% | 5 X 10^6 cells/ml | 1 | 1.5mM cysteine |
| FS025 | 1SN12 | Starch | No | 12 | 5 to 6 | 37C | Anaerobic conditions | NA | 1.0% | NA | 1 | 1.5mM cysteine |
| FS026 | 1SY12 | Starch | Yes | 12 | 4 to 5 | 37C | Anaerobic conditions | Positive | 1.0% | 5 X 10^6 cells/ml | 1 | 1.5mM cysteine |
| FS027 | 2SN12 | Starch | No | 12 | 5 to 6 | 37C | Anaerobic conditions | NA | 1.0% | NA | 1 | 1.5mM cysteine |
| FS028 | 2SY12 | Starch | Yes | 12 | 4 to 5 | 37C | Anaerobic conditions | Positive | 1.0% | 5 X 10^6 cells/ml | 1 | 1.5mM cysteine |
| FS029 | 1BN12_2 | NA | No | 12 | 7 | 37C | Anaerobic conditions | NA | NA | NA | 2 | 1.5mM cysteine |
| FS030 | 1GSY12_2 | Glucose + Starch | Yes | 12 | 6 | 37C | Anaerobic conditions | Positive | 1.0% | 5 X 10^6 cells/ml | 2 | 1.5mM cysteine |
| FS031 | 1BY12_2 | NA | Yes | 12 | 7 | 37C | Anaerobic conditions | Positive | NA | 5 X 10^6 cells/ml | 2 | 1.5mM cysteine |
| FS032 | 2BN12_2 | NA | No | 12 | 7 | 37C | Anaerobic conditions | NA | NA | NA | 2 | 1.5mM cysteine |
| FS033 | 2GSY12_2 | Glucose + Starch | Yes | 12 | 6 | 37C | Anaerobic conditions | Positive | 1.0% | 5 X 10^6 cells/ml | 2 | 1.5mM cysteine |
| FS034 | 2BY12_2 | NA | Yes | 12 | 7 | 37C | Anaerobic conditions | Positive | NA | 5 X 10^6 cells/ml | 2 | 1.5mM cysteine |

**Supplementary Table 3** – Details of all the different conditions and treatments of the samples.
