## Supplementary Table 4 for "Gut microbiota modulation in response to combination of *Escherichia coli* Nissle 1917 and sugars: Lessons from comparative analysis of fecal microbiota of two healthy donors from 2019-2021"

| Sample | Sample ID | Raw Reads | Filtered Reads |
|---|---|---|---|
| FS001 | 1BN0 | 1159700 | 254633 |
| FS002 | 1BY0 | 1071680 | 238609 |
| FS003 | 2BN0 | 992255 | 244484 |
| FS004 | 2BY0 | 934604 | 218251 |
| FS005 | 1BN12 | 1220605 | 265205 |
| FS006 | 1BY12 | 979075 | 208020 |
| FS007 | 2BN12 | 955120 | 225763 |
| FS008 | 2BY12 | 1003250 | 234239 |
| FS009 | 1GSN12 | 1066680 | 235486 |
| FS010 | 1GSY12 | 1171562 | 247516 |
| FS011 | 2GSN12 | 1002200 | 232899 |
| FS012 | 2GSY12 | 997460 | 223082 |
| FS013 | 1GN12 | 1160946 | 257792 |
| FS014 | 1GY12 | 1178518 | 262896 |
| FS015 | 2GN12 | 1077576 | 251102 |
| FS016 | 2GY12 | 1025440 | 231670 |
| FS017 | 1GaSN12 | 1128388 | 253155 |
| FS018 | 1GaSY12 | 1189029 | 257407 |
| FS019 | 2GaSN12 | 1052837 | 268351 |
| FS020 | 2GaSY12 | 918035 | 217708 |
| FS021 | 1GaN12 | 1112326 | 246335 |
| FS022 | 1GaY12 | 1109789 | 243469 |
| FS023 | 2GaN12 | 877441 | 200187 |
| FS024 | 2GaY12 | 892529 | 188982 |
| FS025 | 1SN12 | 1346280 | 297487 |
| FS026 | 1SY12 | 1008663 | 212310 |
| FS027 | 2SN12 | 997096 | 236162 |
| FS028 | 2SY12 | 1009355 | 214536 |
| FS029 | 1BN12_2 | 175956 | 62760 |
| FS030 | 1GSY12_2 | 225978 | 73328 |
| FS031 | 1BY12_2 | 226220 | 77926 |
| FS032 | 2BN12_2 | 163518 | 56455 |
| FS033 | 2GSY12_2 | 193727 | 66172 |
| FS034 | 2BY12_2 | 290575 | 100219 |

**Supplementary Table 4** – Sample-wise counts of the raw reads and filtered reads (Filter method is described in the method section)
